# Microbial ecological insights from estuarine sediments revealed by temporal and spatial network analysis

**DOI:** 10.64898/2026.09.16.752004

**Authors:** Leire Garate, Anthony Chariton, Ion L. Abad-Recio, Anders Lanzén

## Abstract

Association network analyses performed on microbial abundance data provides information regarding ecosystems functioning, quality and potential keystone taxa. Samples for network analyses can be collected spatially, temporally or both. Here, we compare the microbiomes and the reconstructed association networks resulting from temporal *vs* spatial sampling regimes from the Oka estuary in the Spanish Basque Country. The taxonomic composition although similar at Phylum level, diverged more at lower taxonomic levels such as Family. The dissimilarity of spatial replicates was higher than the temporal replicates; however, the dispersion of OTU abundances across samples was similar in the two datasets, indicating that OTU abundances varied similarly across samples. Moreover, the topological properties and keystones differed between networks. The number of edges and the connectance were higher in the spatial network, and keystones were more taxonomically diverse in the temporal network. Among those correlations present in the spatial and temporal networks, we identified potential interactions involved in the sulphur and nitrogen cycles, and in the degradation of organic matter. We conclude that network analyses reveal ecological aspects of the ecosystems functioning, nevertheless, the sampling must be thoroughly planned according to the aim of the study, since different sampling strategies may result in different outcomes.

## INTRODUCTION

Microbial communities form the base of biogeochemical cycles and the trophic webs in ecosystems, and together with their high functional diversity and rapid response to environmental changes, they are a key component in the study of ecosystems health (Guo et al., 2019; Simonin et al., 2019). Previous studies in sedimentary microbiomes from estuaries identified indicators of ecological quality (e.g. Clark et al., 2020; Lanzén et al., 2020). Besides changes in microbial composition and abundance, understanding how the relationships between microorganisms are affected by changes in the environment can provide deeper insights into their ecological functioning.

Association (co-occurrence) networks, based on relative abundances of taxa across the studied communities, provide ecological insights such as the potential interactions between organisms, and can reveal which taxa are likely to be important to community resilience, also named keystone species (Banerjee et al., 2018). This is especially relevant for microorganisms, whose ecological roles and positions in food-webs are often unknown. An increasing number of studies have identified potential keystones based on degree centrality (number of significant correlations established by a taxon) (Brauner et al., 2025; Garate et al., 2025; Iram et al., 2025), although these results should be interpreted with caution as they may vary depending on parameters such as the data preprocessing or the tool used to calculate the network. On the other hand, topological metrics obtained from these networks can indicate the ecological state of environments, being also useful resources for evaluating ecosystems (Codello et al., 2022). For example, higher values of modularity may appear in sites submitted to some impact, and a higher proportion of negative correlations may be linked to a greater stability of the community (Herren & McMahon, 2018; Garate et al., 2025).

It is recommended that such network analyses use as many samples as possible, and at least 20-25, (Berry & Widder, 2014). These samples can be collected across a temporal series, spatially, or both, i.e. using spatial replicates, repeated in a time series. This represents a fundamental decision in terms of sampling strategy, and while studies focused on microbial ecology differ in such strategies, there are few or no published studies that have compared the consequences of this choice, i.e. compare how networks based on spatial *vs* temporal samples may vary. To this end, we compared samples collected across a temporal series with samples across a spatial grid, from the same sampling site in the Oka estuary (Basque Country, Spain), and used them to calculate, analyse and compare the resulting prokaryotic community association networks, based on eDNA metabarcoding.

## MATERIAL & METHODS

### Samples collection

Samples were collected by scraping the top centimetre of the sandy, intertidal estuary sediments into a sterile 50 ml Falcon tube and were kept on ice in a portable cooler until arrival to the laboratory.

Two sampling strategies were carried out. Firstly, sediment samples from temporal series were collected from the same exact location (approximately 3 × 3 m) yearly or twice yearly from 2015-2019 and complemented with more frequent sampling, twice monthly, from January 2020 to January 2021, collecting a total of 28 samples. In addition, spatial sediment sampling was carried out in June 2021 from a grid surrounding the temporal sampling site, from three parallel lines separated by 5 meters. Each line was sampled at one-meter intervals with 10 samples from each line, resulting in a total of 30 samples. One of the spatial replicates was removed from the study because the number of reads obtained from it was lower than 5000. All the samples were kept frozen (−80°C) until eDNA extraction.

### Nucleic acid extraction and metabarcoding

Total DNA was extracted using DNeasy PowerSoil Pro Kit (Qiagen) following the manufacturer’s instructions. Nucleic acid extracts were visualised on an electrophoresis gel, and their concentration and quality were measured with a Nanodrop spectrophotometer and Qubit dsDNA BR Assay Kit, used in a Qubit fluorometer. Sample concentrations were normalised to 10 ng/µl for 16S rRNA amplification.

The V4 region of the 16S small subunit rRNA gene was amplified with primers 519F (CAGCMGCCGCGGTAA; Øvreås et al., 1997) and 806RD (GGACTACNVGGGTWTCTAAT; Apprill et al., 2015) as described previously by Lanzén et al.(2020). Two negative controls per plate were added before DNA extraction and PCR. Samples were amplified as follows: 95 °C for 3 minutes, 30 cycles of 95 °C for 30 s, 50 °C for 30 s and 72 °C for 45 s, and a final extension at 72 °C for 5 minutes. PCR triplicates were then pooled and purified using AMPure XP beads (Beckman Coulter). 5 µl of purified amplicons were used as template for a second PCR round in which Illumina Nextera adapters were attached to the amplicons. The program used in the second PCR was: 95 °C for 3 minutes, 8 cycles of 95 °C for 30 s, 55 °C for 30 s, and 72 °C for 30 s, followed by a final extension at 72 °C for 5 min. The second PCR product was again purified using AMPure XP beads and eDNA concentration was measured using a Qubit fluorometer. PCR products yielding sufficient amount and quality of amplified DNA were then pooled in into a final library at equimolar concentrations of 10 nM. Products yielding insufficient quantities for this pooling (negative controls) were added columetrically 5 µl. Sequencing was carried out using Illumina MiSeq with paired-end 2 × 300 bp v3 chemistry, carried out at the National Genome Analysis Centre (CNAG - Centre Nacional d’Anàlisi Genòmica, Barcelona) and distributed over four different runs. Raw sequences are available from the INSDC Sequence Read Archive with BioProject accession number PRJEB124988.

### Sequence analyses

Sequence data processing was performed as described previously (Lanzén et al. 2020). Read-pair overlapping and sequence data filtering was carried out using vsearch v2.7.1(Rognes et al., 2016) allowing 20 mismatches, and primers removed using cutadapt v1.18(Martin, 2011), followed by truncation to 252 bp, discarding reads without full and correct primer sequences or more than one expected error. The remaining sequences were de-replicated and sorted by abundance using *vsearch* and clustered into sequence variants (SVs) using *SWARM* v2.21 (Mahé et al., 2015). Abundances of unique sequences across samples were retained to construct an SV contingency table, using the scripts *fasta_merging*.*py* and *matrix_creation*.*py*, of *SLIM* (Dufresne et al., 2019). We then discarded SVs with a total abundance across samples of one (singletons), then putative chimeras, using *vsearch*, reference based with SilvaModPR2 v138 (Lanzén et al., 2012; https://github.com/lanzen/CREST) followed by de novo mode. To correct remaining PCR and sequencing artefacts and to merge intra-specific or intra-genomic SVs, we then applied LULU post-clustering curation (Frøslev et al., 2017) with default parameters except for increasing minimum similarity to 97%). Curated SVs were aligned to SilvaModPR2 v138 using *blastn* v2.6.0+ and then taxonomically classified using *CREST* v3.1.0 (Lanzén et al. 2012).

Unclassified SVs at kingdom rank and non-target SVs originating from Eukaryota (18S or organellar SSU; n=5651) were then removed along with 47 putative (p<0.05) contaminant SVs in R software (v4.6.1; R Core Team, 2021), identified based on PCR and extraction blanks, with *decontam* R package v.1.12.0 (Davis et al., 2018). Cross-contamination was reduced by setting OTU abundances to zero where SVs occurred in a sample at an abundance being 100x lower relative to its average non-zero abundance across samples (corresponding to the UNCROSS algorithm, Edgar, 2016). To decrease bias from uneven sequencing depth across samples, all rare SVs defined as those that never reached 0.1% abundance in at least one sample, were removed.

Filtered SVs with identical taxonomic annotation (and their abundances across time-point samples) were merged into operational taxonomic units (OTUs) based on taxonomy, for reconstruction of association networks. This was done by merging all SVs with identical annotations, without a fixed taxonomic rank, meaning that our final list includes both species rank OTUs (e.g. *Desulfuromusa bakii*) as well as “orphan” OTUs for the parent genus or higher ranks summarising all SVs unclassified at species or lower ranks (e.g., *Arenimonas*, Sedimenticolaceae.).

### Network reconstruction and visualisation

Ecological association networks were calculated with *CCREPE* R package (Schwager et al. 2014) using the *ccrepe* function, and filtering those correlations with *q value*s <0.01. In order to have comparable networks calculated with the same number of samples, we used bootstrapping, randomly selecting 28 samples at each iteration reconstructing an association network from the spatial replicates, a total of 100 times, to generate 100 different networks. In the final spatial network, we retained all edges (correlations) that consistently showed the same sign and appeared in 90% of the individual networks. The resulting networks were exported to Cytoscape v.3.9.1 (Shannon et al. 2003) for visualization and topological analyses with Cytoscape tool Analyse Network. Networks were also analysed using the R package *igraph* (Csardi and Nepusz 2006). Potential keystone taxa were defined as previously described by Garate et al. (2022), selecting the top ten nodes with the highest sums of their degree of connectivity (number of connections to different nodes) and closeness centrality (how close a node is to all others). Finally, we compared the temporal and spatial networks to find the common correlations retrieved by the two sampling regimes.

## RESULTS & DISCUSSION

### Differences and similarities across microbial communities

Taxonomically, microbial communities collected temporally and spatially were relatively similar, with Proteobacteria being the most abundant phylum, and unclassified Gammaprotebacteria the most abundant taxon across ranks (Figure 1 A). One of the major differences between the datasets is the higher abundance of Desulfobacterota in the time series communities. More differences can be observed at family level, where families that showed higher abundances in winter-early spring from the temporal samples were absent in the spatial replicates (Figure 1 B).

**Figure 1.**
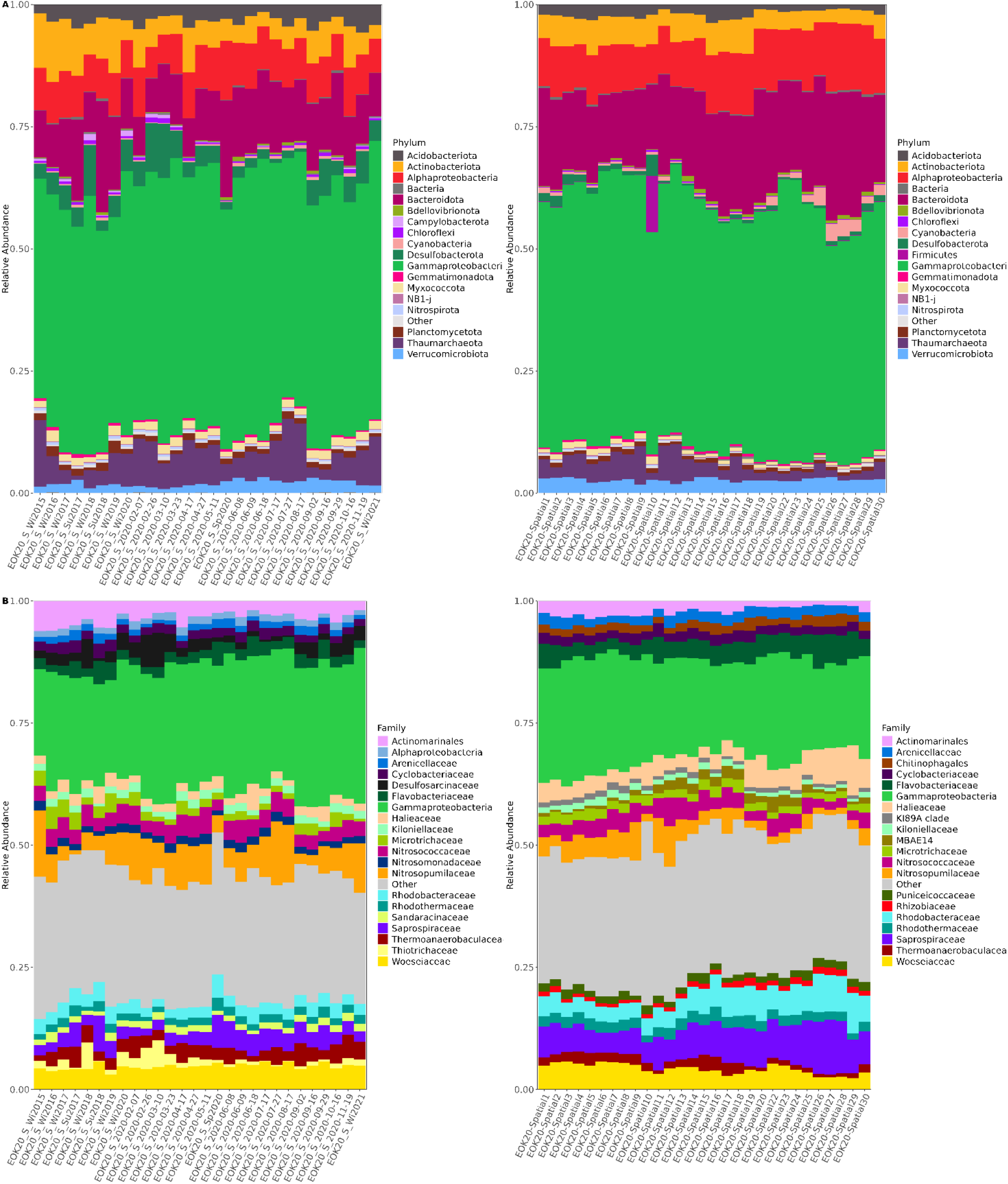
Composition plots of the relative abundance of the 20 most abundant phyla (A) and families (B) across the temporal (left) and spatial (right) replicates.

Pairwise Bray-Curtis dissimilarities among samples in the same dataset (temporal and spatial) differed significantly between sampling strategies, with spatial replicates showing a significantly higher median dissimilarity (Figure 2 A-C). Pairwise dissimilarity increased with and was best explained by seasonal difference (days of the year apart, disregarding sampling year) and spatial difference in meters, respectively (modelled using a non-linear distance-decay relationship with least-squares regression and a linear relationship, respectively), showing similar dispersion. However, the coefficient of variance (standard deviation of OTU abundances divided by their means; Figure 2 D), which instead measures how dispersed OTU abundances are across samples, did not differ significantly between the temporal and spatial dataset. These results indicate that the microbial communities collected in the spatial replicates from a relatively small area (10 × 15m) differed more compared to those collected from the same location through time.

**Figure 2.**
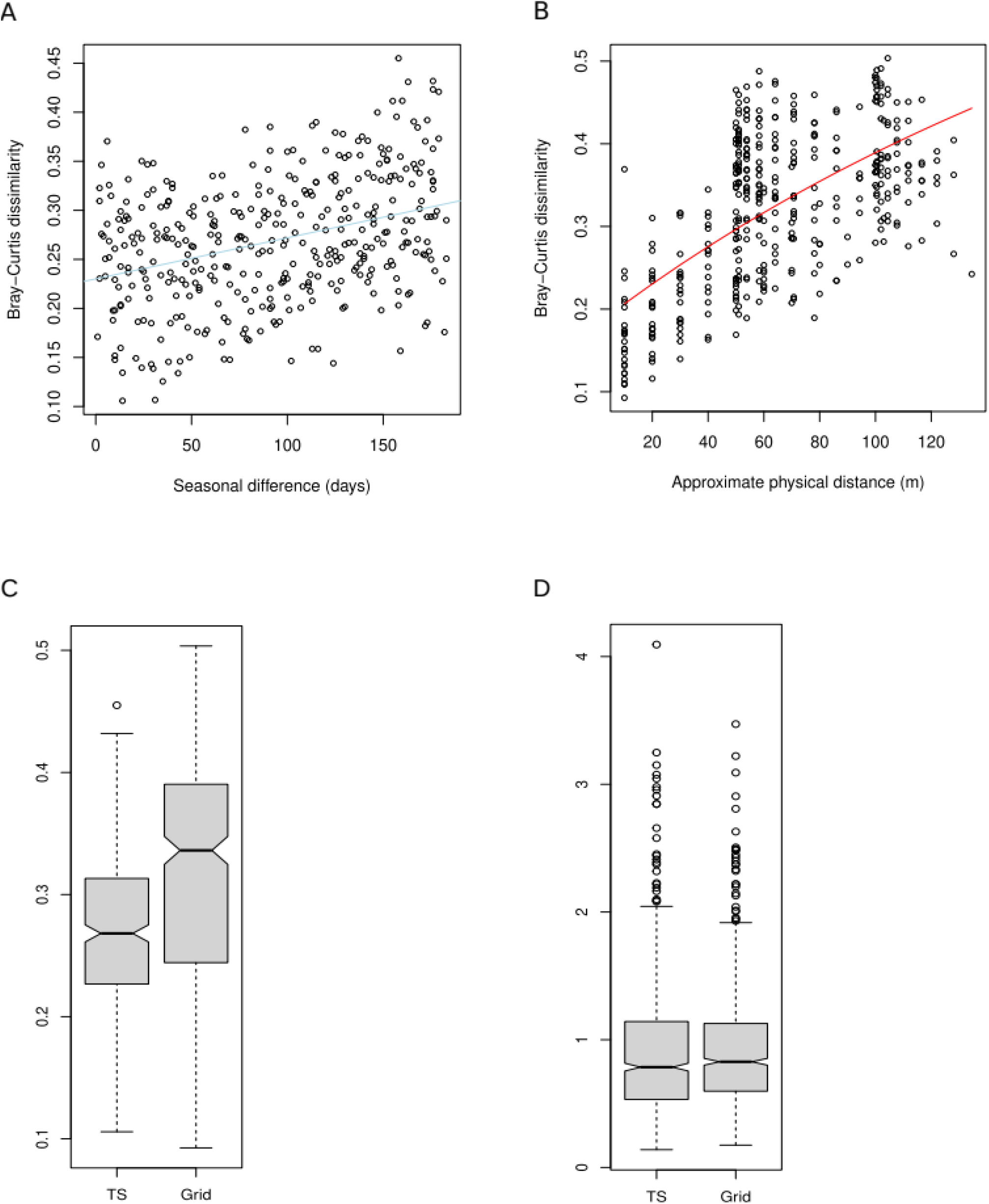
Bray-Curtis dissimilarities among samples in the temporal (A) and spatial (B) sampling strategies. (C) Pairwise median dissimilarity in the temporal series (TS) and the spatial sampling (Grid). (D) Coefficient of variance (standard deviation of OTU abundances divided by their means) in the temporal series (TS) and spatial sampling (Grid).

### Temporal and spatial association networks and common correlations between them

Although the networks were calculated using the same parameters, number of samples and applying the same prevalence filtering, temporal and spatial networks differed notably in terms of topology and keystone taxa identification (Supplementary Figure 1). Table 1 shows the topological properties of the two networks. The clearest difference is the elevated number of edges in the spatially derived network, resulting also in a higher value of connectance. Moreover, the identified keystone taxa differed between networks (Table 2). When reconstructing a consensus network limited to edges (correlations) found in both networks, we identified a total of 91 associations between 50 different taxa (Figure 3). Based on bibliography, the taxa present in each of the modules retrieved had a metabolic potential that made it possible to classify these modules in two microbial biogeochemical processes: the sulphur cycle (63 associations) and nitrification (6 associations). There are also several modules composed of taxa that are potentially involved in degradation of organic matter (22 associations). Interestingly, nine of the top ten keystones from the temporal network appeared in the cluster purportedly involved in sulphur metabolism, being mainly anaerobic taxa. The presence of bacteria involved in sulphur metabolism in the surface sediments has previously been reported in undisturbed temperate and tropical estuaries (Garate et al., 2025; Shah et al., 2021). This suggests that they play a more important role in oxygenated marine sediments than what is sometimes assumed, (Bühring et al., 2005). Hence, they likely influence the carbon and sulphur cycles via dissimilatory sulphur reduction, (Caffrey & Voordouw, 2010; Le Gall & Postgate, 1973; Stoeva & Coates, 2019).

**Table 1.**
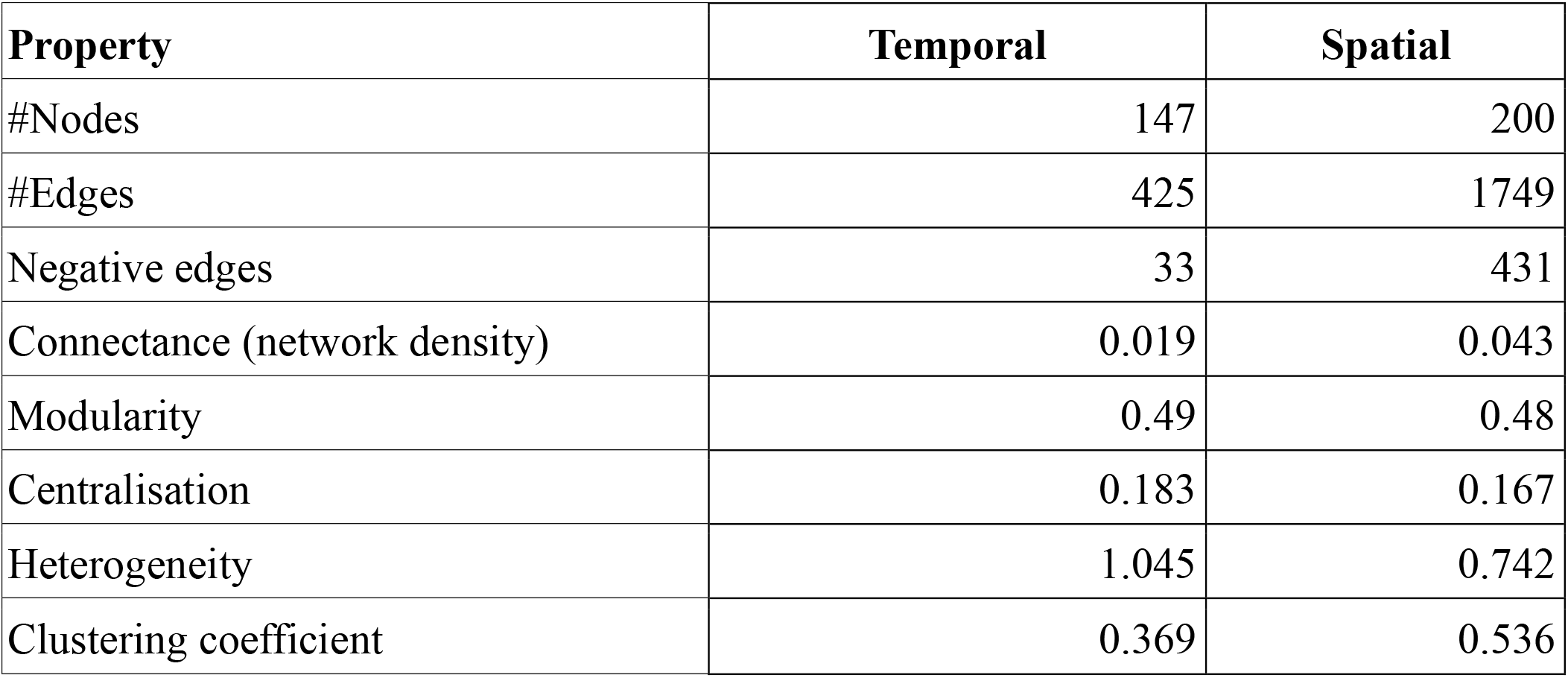
Topological properties of temporal and spatial networks.

| Property | Temporal | Spatial |
| --- | --- | --- |
| #Nodes | 147 | 200 |
| #Edges | 425 | 1749 |
| Negative edges | 33 | 431 |
| Connectance (network density) | 0.019 | 0.043 |
| Modularity | 0.49 | 0.48 |
| Centralisation | 0.183 | 0.167 |
| Heterogeneity | 1.045 | 0.742 |
| Clustering coefficient | 0.369 | 0.536 |

**Table 2.** Keystone taxa and Classes identified in temporal and spatial networks. In bold are the taxa found in consensus networks built from the individual networks.

| Temporal Network |  | Spatial Network |  |
| --- | --- | --- | --- |
| Taxon | Class | Taxon | Class |
| **Desulfosarcinaceae** | Desulfobacteria | Hyphomonadaceae | Alphaproteobacteria |
| **Desulfobulbaceae** | Desulfobulbia | **Flavobacteriaceae** | Bacteroidia |
| **SEEP-SRB1** | Desulfobacteria | **Lutimonas** | Bacteroidia |
| **Anaerolineaceae** | Anaerolineae | **Rhodobacteraceae** | Alphaproteobacteria |
| **B2M28** | Gammaproteobacteria | **Alphaproteobacteria** | Alphaproteobacteria |
| **Sva0081 sediment group** | Desulfobacteria | **Roseobacter** | Alphaproteobacteria |
| **Thiotrichaceae** | Gammaproteobacteria | Thiohalorhabdaceae | Gammaproteobacteria |
| **Calditrichaceae** | Calditrichia | **Actibacter** | Bacteroidia |
| Bacteroidetes BD2-2 | Bacteroidia | Ilumatobacter | Acidimicrobiia |
| **Sva0485** | Sva0485 | **Halioglobus** | Gammaproteobacteria |

**Figure 3.**
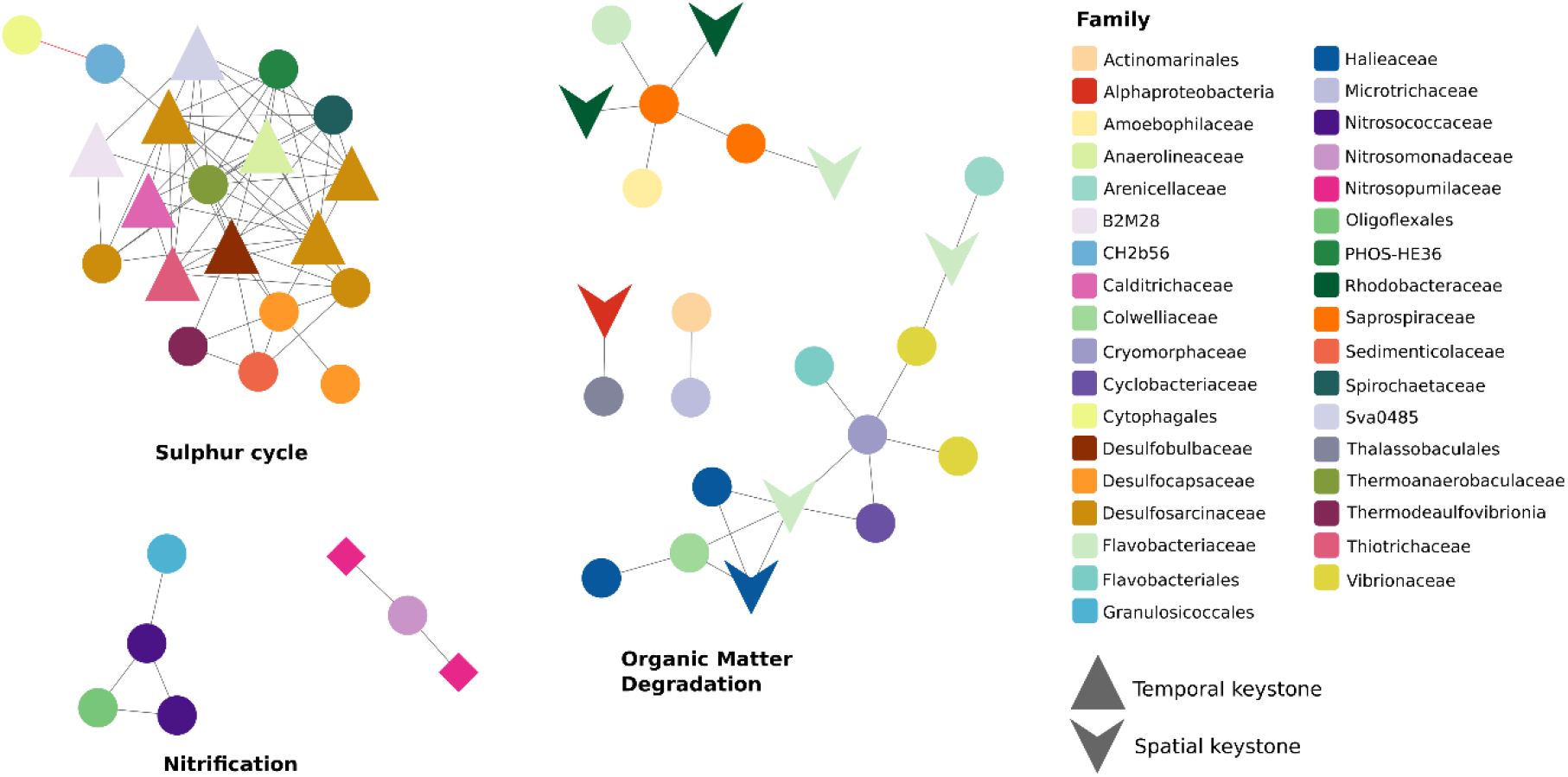
Common associations that are present in the temporal and the spatial networks. Nodes represent taxa colored by family. Grey lines correspond to positive correlations, while the red line corresponds to the negative correlation retrieved in both networks.

On the other hand, seven of the top ten keystones identified in the spatially based network appeared to be involved in organic matter degradation. This included, three taxa belonging to the family Flavobacteriaceae, and two to the Rhodobacteraceae family (Kirchman, 2006; Rajeev & Cho, 2024).

Some edges of taxa involved in nitrification were identified in the two networks. For example, a taxon from *Nitrosomonas* (Betaproteobacteria) correlated with two from the Nitrosopumilaceae (Thaumarcheaeota). Oxidation of ammonium can be carried out both by ammonia-oxidizing bacteria and archaea, and potentially the amount of ammonium in the environment is responsible for the niche differentiation of ammonia oxidizers (French et al., 2021).

Several of these keystone taxa in both the temporal and spatial networks have been previously identified in studies as indicators of environmental quality. For example, taxa associated with Hyphomonadaceae and Halioglobus, were identified as keystones and connectors (those that presented high betweenness centrality) in Basque estuaries that presented good ecological status (Garate et al., 2025). Although the Oka estuary has been classified as presenting good ecological quality (Basque Monitoring Network; Borja et al., 2022), there have also been identified taxa previously reported as indicators of poor sediment quality (Lanzén et al., 2020), namely Bacteroidetes BD2-2 and Anaerolineaceae, as well as the previously mentioned taxa associated with sulphur metabolism, i.e. Thiotrichaceae and Desulfobulbaceae.

The only negative correlation common to both networks appeared in the module involved in sulphur metabolism and took place between a member of the aerobic Cytophagales and a taxon from Gammaproteobacteria class “CH2b56”, which is likely anaerobic (Flood et al., 2021; Shaner, 2022)).

In contrast, in the temporal series, these two taxa followed an opposite trend over time (Supplementary Figure 2), suggesting that they may react in opposite manners to the temporal trends in oxygenation.

In conclusion, this study showcased how the type of sampling can provide different results of network analyses, despite the samples being collected and treated equally. This is an important issue to consider depending on the questions the researchers are seeking answers to. Regardless of the discrepancies observed between the spatial and temporal networks, we revealed some of the metabolic processes that potentially occur in estuarine sediments, such as those related with sulfur and nitrogen metabolism, or the organic matter degradation. Hence, our findings emphasize the applicability of ecological networks in improving our knowledge of the smallest components of these ecosystems, and their functioning.

## ACKNOWLEDGEMENTS

We would like to thank Iñaki Mendibil for his assistance in the eDNA isolation, amplification, and library preparation. This is contribution number [to be filled in at acceptance] from the Marine Research Division of AZTI.

